# A root pathogen drives rhizosphere enrichment of antagonistic *Pseudomonas* and induces production of an antimicrobial metabolite

**DOI:** 10.64898/2026.08.26.747274

**Authors:** Mohamed Zouaoui, Marie Solau, Aurélien Amiel, Alice Penrose, Jean Belleville, Quentin Bazerque, Sylvie Fournier, Amélie Perez, Laurent Camborde, Elodie Gaulin, Thomas Rey, Bernard Dumas

## Abstract

In plants, the development of soil-borne diseases has been shown to trigger the recruitment of beneficial microbes, which contribute to defense against pathogens. However, the underlying mechanisms driving this recruitment remain elusive. Here, we used a gnotobiotic system combined with a synthetic bacterial community (SynCom) derived from the *Medicago truncatula* rhizosphere to dissect microbiota–pathogen interaction and its role in the development of root rot caused by *Aphanomyces euteiches*, a devastating soilborne oomycete pathogen of legumes. Through integrated metabarcoding, metabolomics and transcriptomics, we reveal that pathogen infection restructures the bacterial SynCom, selectively enriching the microbial community with specific *Pseudomonas* spp. strains displaying anti-*A. euteiches* activity. This shift alleviates root rot symptoms, triggers the biosynthesis of the antibiotic 2,4-diacetylphloroglucinol (DAPG), a *Pseudomonas* specialized metabolite inhibiting *A. euteiches* growth, and amplifies the plant’s endogenous isoflavonoid defense responses. Unexpectedly, we found that *A. euteiches* directly activates DAPG production in beneficial bacteria independently of the plant, through the production of a heat-stable, high-molecular-weight (30–100 kDa) extracellular components. These findings uncover a novel mechanism whereby a pathogen inadvertently activates antibiotic production in beneficial bacteria, extending the plant immune system. Our research underscores the critical role of microbial interactions in the rhizosphere in determining root disease outcomes, paving the way for microbiome-based strategies to combat root diseases.

## Introduction

The plant rhizosphere is colonized by a highly diverse and abundant microbial consortium that plays crucial roles in plant nutrition and resistance to both biotic and abiotic stresses. Numerous studies have demonstrated that the assembly of the root microbiota is partially driven by the plant itself and can adapt in response to environmental changes, particularly under biotic stress, favoring the enrichment of beneficial species [1, 2].

One of the most compelling examples of plant–soil–microbe interactions demonstrating adaptation to pathogenic environments is the establishment of disease-suppressive soils. These are defined as soils in which the incidence or severity of plant disease remains low despite the presence of a virulent pathogen, a susceptible host, and conducive environmental conditions [1]. This phenomenon has been documented in numerous agroecosystems and for various soil-borne diseases [3]. Disease suppressiveness can be classified as either broad-spectrum (general) or pathogen-specific. General suppressiveness arises from the collective activity of the soil microbiome, where a diverse and competitive microbial community limits pathogen proliferation through mechanisms such as nutrient competition, antibiosis, or parasitism. In contrast, specific suppressiveness is associated with particular microbial taxa or consortia that directly antagonize pathogens. A well-documented example of specific suppressiveness involves *Pseudomonas* strains producing 2,4-diacetylphloroglucinol (DAPG), which suppress take-all disease in wheat [4, 5].

An emerging concept from these observations is the "cry for help" hypothesis, which proposes that plants under attack actively recruit beneficial microbes to their rhizosphere to enhance resistance [6]. Mechanistically, this process could be mediated by root exudates— which include sugars, amino acids, organic acids, and a wide array of secondary metabolites—that serve as chemical signals for soil microorganisms [7, 8]. Pathogen infection has been shown to alter both the quantity and composition of root exudates, leading to the secretion of specific metabolites that attract antagonistic bacteria [2, 9, 10]. These bacteria, in turn, suppress pathogens through competition, antibiosis, or the induction of systemic resistance [2, 9, 10].

*Aphanomyces euteiches* is a soil-borne oomycete pathogen responsible for root rot in a wide range of leguminous crops, including pea, lentil, alfalfa, and bean, as well as the model legume *Medicago truncatula* [11, 12]. In addition to *A. euteiches*, other pathogenic fungi and oomycetes, such as *Fusarium* and *Pythium* species, contribute to the disease collectively referred to as the ‘root rot complex’ [12, 13]. Conventional control measures, including chemical fungicides and crop rotation, often provide only limited or inconsistent protection, largely due to the broad host range and long-term survival of these pathogens. While breeding for resistance has identified partial resistance genes in legumes, durable resistance remains elusive [14]. Meanwhile, biological control strategies have underscored the role of beneficial microbial species in reducing disease severity [15, 16].

In our recent work, we demonstrated the beneficial activity of the mycoparasitic oomycete *Pythium oligandrum* against *Aphanomyces* root rot, primarily through the induction of plant defense responses such as the production of specific isoflavonoids [17]. This suggests that optimizing the rhizosphere microbiota through the presence of beneficial microbial strains could represent an effective strategy to improve legume resistance to *Aphanomyces* root rot. A complementary strategy could be the selection of legume genotypes capable of recruiting beneficial microorganisms. Recent studies have correlated the abundance of specific microbial groups with enhanced resistance of pea cultivars to pathogens involved in root rot disease [18, 19]. Notably, the presence of some specific microbes in the rhizosphere is correlated with root rot resistance, implying that the composition of the rhizosphere microbiota is at least partially influenced by the plant genotype and could serve as a potential means of resistance against legume root rot [18, 19].

Here, we investigated whether rhizospheric bacteria could influence the outcome of the *M. truncatula–A. euteiches* interaction and the mechanisms underlying this effect. As a first step, we designed a synthetic community (SynCom) comprising selected strains isolated from the *M. truncatula* rhizosphere. These strains were chosen for their demonstrated biological activities related to plant nutrition and development (e.g., phosphate solubilization, nitrogen fixation, and phytohormone production) as well as their protective roles against biotic stress (e.g., anti-*A. euteiches* activity). The SynCom was uses in gnotobiotic experiments coupled with multi-omic analyses, revealing that *A. euteiches* infection selectively enriches DAPG-producing *Pseudomonas* strains in the rhizosphere, and induces DAPG production, a metabolite with anti-*A. euteiches* activity. We provide evidence that bacterial DAPG production is triggered by the perception of high-molecular-weight, heat-stable extracellular compound released by *A. euteiches*.

## Materials and Methods

### Bacterial Isolation and Cultivation

Rhizosphere-associated bacteria were isolated from *Medicago truncatula* F83005.5 grown in potting soil for 2 months using the limit dilution high-throughput cultivation method [20]. Root samples were washed with sterile phosphate-buffered saline (PBS) to recover rhizosphere-associated bacterial cells. The resulting bacterial suspension was centrifuged and filtered through a 45 μm membrane to remove plant debris and larger particles. The filtered bacterial suspension was serially diluted in 10% TSB, and a cumulative dilution of 1:48,600 (obtained through successive dilution steps) was retained such that approximately 30% of wells showed bacterial growth, following Poisson distribution principles to ensure single-cell isolation. Twenty-five 96-well plates were inoculated at this dilution and incubated at 24°C in darkness for 7–10 days in 10% TSB. Wells showing bacterial growth (OD_600_ > 0.1) were subsequently consolidated into 12 pooled 96-well plates for downstream processing.

For routine maintenance, the retained isolates were grown in 10% Tryptic Soy Broth (TSB) in 96-well plates at 24°C in darkness. For solid medium preparation, 15 g L⁻¹ agar was added to create Tryptic Soy Agar (TSA). Following cultivation, sterile glycerol (25% final concentration) was added to liquid cultures for cryopreservation at −80°C. Wells showing bacterial growth were lysed by alkaline lysis combined with a high-temperature treatment and subjected to 16S rRNA gene (V3–V4 region) PCR amplification. Well-specific barcodes (tags) were added to the primers during a first PCR so that each amplicon could be traced back to its plate and well of origin, followed by a second PCR adding the Illumina adapters. PCR products from positive wells were pooled, multiplexed, and sequenced at the GeT-Biopuces platform (Genotoul, Toulouse, France) on an Illumina MiSeq instrument with v2 chemistry, generating 2 × 250-bp reads according to the manufacturer’s instructions.

### *Aphanomyces euteiches* culture and zoospore production

*Aphanomyces euteiches* isolate MF1 was maintained on corn meal agar (CMA, 17 g L⁻¹) at 22°C in darkness. Zoospores were produced as previously described [21] and concentration was determined by counting with a Fuchs-Rosenthal hemocytometer under a standard binocular light microscope.

### 16S rRNA gene processing and OTU assignment of the culture collection

To taxonomically assign each bacterial isolate and confirm purity, the multiplexed 16S rRNA amplicons were demultiplexed and processed using QIIME2 (v2024.2) [22]with VSEARCH for de novo OTU clustering at 97% sequence similarity. Reads were assigned to their well of origin using the dual-index barcodes. Wells in which more than one OTU was detected were considered non-pure and excluded from downstream analysis. Taxonomy was assigned to each representative OTU sequence against the SILVA reference database [23].

### Phosphate solubilization activity

Bacterial isolates were screened for their ability to solubilize inorganic tricalcium phosphate (TCP) and organic phytate using a modified phytase screening medium [24]. The medium contained (per liter): 10 g glucose, 2 g NH₄NO₃, 0.5 g KCl, 0.5 g MgSO₄·7H₂O, 0.01 g MnSO₄·H₂O, 0.01 g FeSO₄·7H₂O, and 20 g agar, adjusted to pH 7.0. For TCP screening, 2 g TCP was added before autoclaving. For phytate screening, filter-sterilized (0.22 μm) phytate solution was added post-autoclaving to achieve a final concentration of 2 g L⁻¹, along with 2 g CaCl₂. Each isolate was spotted individually onto the screening medium, and plates were incubated at 24°C in darkness for 12 days. Phosphate solubilization activity was assessed by the presence of a clear zone (halo) around bacterial colonies and scored in a binary manner as either negative (0, no halo) or positive (1, halo present). To confirm genuine phytate degradation and eliminate false positives arising from medium acidification (halos caused by pH-dependent solubilization rather than phytate hydrolysis), plates were treated with 2% (w/v) cobalt chloride solution for 20 min at room temperature [25]. Disappearance of halos indicated false positive results.

### Auxin production assay

Indole-3-acetic acid (IAA) production was detected using Salkowski’s reagent (12 g FeCl₃ L⁻¹ in 7.9 M H₂SO₄) as described by [26]. Bacterial cultures in 96-well plates were treated with 100 μL reagent per well. Pink coloration indicated IAA presence (detection limit: 6 μg mL⁻¹). A standard curve was prepared with IAA concentrations ranging from 1.56 to 100 μg mL⁻¹ in seven dilution steps.

### Nitrogen fixation screening

Diazotrophic potential was assessed by PCR amplification of the *nifH* gene using primers Ueda19F/R6 [27]. DNA was extracted from bacterial cultures as described for the cultivation step above. PCR products were analyzed using qPCR with SYBR Green (2 μL PCR product, 10 μL SYBR Green 1× in 384-well plates) on a BioRad CFX real-time PCR system. Four confirmed diazotrophic strains were used as positive controls: FSM-MA (alfalfa), USDA110 (soybean), 1AE200 (lupin), and R7A (lotus).

### Detection of anti-oomycete activity

Bacterial isolates were grown on potato dextrose agar (PDA) for 7 days, then co-cultured with *A. euteiches* zoospores (2,500 zoospores mL⁻¹) by spray inoculation for a preliminary screening. Plates were incubated at 22°C and examined at 6- and 11-days post-inoculation for mycelium growth inhibition. Candidate strains were then confirmed in a dual confrontation assay. Bacteria were streak-inoculated on one third of a PDA plate and incubated at 22°C in darkness for 3 days. An *A. euteiches* mycelium plug was placed on the opposite side of the plate. Inhibition zones were measured using ImageJ 1.54d at 9- and 11-days post-inoculation and expressed as percentage of mycelial growth relative to the no-bacteria control.

To assess the contribution of volatile organic compounds (VOCs) to *A. euteiches* inhibition, selected *Pseudomonas* strains were co-cultured with *A. euteiches* mycelium plugs on a compartmentalized Petri dish (PDA, 39 g L⁻¹) that physically separated bacteria from the pathogen, preventing direct contact while allowing volatile exchange. Plates were incubated at 22°C in darkness for 4 days. To assess the contribution of siderophore-mediated iron competition, *Pseudomonas* strains were co-cultured with *A. euteiches* on PDA supplemented with FeCl₃ (0.13 g L^-1^) to saturate siderophore activity and eliminate iron competition as a mechanism. Plates were incubated at 22°C in darkness for 4 days.

Anti-*Aphanomyces* activity of specialized metabolites (DAPG, MAPG and PG) was assessed in 96-well microplates (200 µL). Each well contained 100 µL of PDB, 50 µL of *A. euteiches* MF1 zoospore suspension (10,000 zoospores mL⁻¹ in sterile Volvic water), and 8 µL of the compound at the desired concentration. Mycelial growth was evaluated by measuring optical density at 600 nm (OD_600_) using a Victor Nivo plate reader (PerkinElmer) immediately after inoculation and after 4 days of incubation at 22°C in darkness (n = 6 replicates per condition).

### SynCom assembly

From the 79 OTUs retained after quality filtering, a SynCom was assembled by selecting strains prioritizing taxonomic representativeness and functional complementarity. Each selected strain was individually cultivated on TSA (10%) plates and transferred into TSB (10%) liquid medium. All cultures were adjusted to an optical density (OD_600_) of 0.1, then pooled in equal volumes (1 mL per strain) to assemble the SynCom. The resulting consortium was stored as glycerol stocks at −80°C at a final concentration of approximately 3.5 × 10⁷ CFU mL⁻¹. For inoculation, the SynCom stock was diluted in half-strength MS medium to a final concentration of 10⁴ CFU per flowpot.

### Flowpot assays

The flowpot system consisted of transparent plastic boxes, each containing six flowpots arranged in a 2×3 configuration on parallel metallic rails [28]. Each flowpot was filled with sterile zeolite substrate (60% particle size 1-2.5 mm; 40% particle size 0.5-1 mm) and equipped with a metallic filter (100 μm pore size) at the base to prevent root leakage, with a 5 mL glass Petri dish positioned underneath to collect flow-through solutions. Seeds of the highly susceptible line *M. truncatula* F83005.5 [29] were surface-sterilized by immersion in concentrated sulphuric acid (H₂SO₄) for 5 min, rinsed three times with sterile water, then immersed in 3% sodium hypochlorite solution for 2 min, and rinsed again three times with sterile water. Seeds were then placed on 0.9% water-agar plates and vernalized at 4°C in darkness for 5 days to synchronize germination. Germinated *M. truncatula* seeds were sown in each flowpot and immediately watered with 10 mL of half-strength Murashige and Skoog (MS) medium (2.2 g L⁻¹) containing the synthetic community (SynCom) which each strain adjusted to OD 0.1. Plants were grown in a phytotron under a 16:8 h light:dark photoperiod at 22°C. After 3 days of plant establishment, half of the flowpot boxes were inoculated with 5 mL of *A. euteiches* zoospore suspension (5,000 zoospores mL⁻¹ in sterile Volvic water), while control flowpots received 5 mL of sterile Volvic water. Following pathogen inoculation, flowpot boxes were returned to the phytotron and maintained under the same conditions, with weekly watering using 5 mL of MS medium for three weeks. Each condition comprised six biological replicates; each biological replicate consisted of material pooled from two plants.

### Pot experiments

Pot experiments were conducted under non-sterile conditions in a phytotron. Plastic pots (8.7 cm diameter, ∼300 mL capacity, autoclavable) were filled entirely with vermiculite and supplemented with 100 mL of one-tenth strength Murashige and Skoog medium (MS 1/10, 0.22 g L⁻¹). Each pot was placed inside a Magenta™ GA-7 vessel (C0542, Merck, Darmstadt, Germany) box to allow bottom-up watering. The assembled system was autoclaved (121°C, 20 min) prior to use. Surface-sterilized and vernalized *M. truncatula* F83005.5 seeds were sown in each pot. Plants were inoculated at sowing with either the complete SynCom or a DAPG-depleted SynCom lacking these strains (SynC⁻), prepared as described for the flowpot assay. Plants were grown in a phytotron under a 16:8 h light:dark photoperiod at 22°C and watered once per week with 50 mL of sterile osmotic water. After 3 days of plant establishment, half of the pots were inoculated with 5 mL of *A. euteiches* zoospore suspension (5,000 zoospores mL⁻¹ in sterile Volvic water); control pots received 5 mL of sterile Volvic water. Plants were harvested 21 days post-inoculation. Root length was measured using ImageJ (v1.54f), tracing the primary root of each plant with the segmented line tool and converting the measured length to centimeters against a scale reference included in each scan. Total DNA was extracted from ground root tissue using the ZymoBIOMICS DNA Miniprep Kit (Zymo Research) following the manufacturer’s protocol.

Detection of the *phlD* gene, encoding the key enzyme of DAPG biosynthesis was performed on genomic DNA of each SynCom strain with the primers phlDCompF1 (5′-ACTTTATTGGCTTCTCGCCG-3′) and phlDCompR1 (5′-GGGTGTCGGTGTTTTATCGGT-3′), designed to amplify a 1,104 bp fragment of the *phlD* gene [30]. Amplification products were resolved by electrophoresis on a 1.5% agarose gel and visualized under UV light, using the DAPG-producing strains A05H and D08H as positive controls.

### 16S rRNA metabarcoding

Rhizosphere samples were collected 21 days post-inoculation. DNA was extracted using the ZymoBIOMICS DNA Miniprep Kit (Zymo Research) following the standard protocol. Full-length 16S rRNA genes were amplified using GoTaq G2 Hot Start Polymerase (Promega) with the Oxford Nanopore 16S Barcoding Kit 1–24 (SQK-16S024 protocol; Oxford Nanopore, 2019). Each barcoded library was cleaned using SparQ PureMag beads and adjusted to 5 ng µL⁻¹. Libraries were pooled by adding 1 µL of each barcoded library to a common DNA LoBind tube (Eppendorf). The pooled library was loaded onto a primed flow cell and sequenced for 18 h on a MinION device controlled by MinKNOW (v24.02). Reads were processed using the NanoClust pipeline [31]. Community composition analyses (alpha diversity, PCoA on Bray-Curtis dissimilarity, PERMANOVA) and differential abundance testing (DESeq2) were performed in R using the phyloseq and DESeq2 packages.

### Genome sequencing and analysis

Genomic DNA from *Pseudomonas* strains was extracted using the Wizard® Genomic DNA Purification Kit (Promega) following the manufacturer’s protocol. Sequencing libraries were prepared using the Rapid Barcoding Kit 96 V14 (SQB-RBK114-96, Oxford Nanopore Technologies) following the manufacturer’s protocol. Sequencing was performed on an Oxford Nanopore MinION using an R10.4 flow cell. Data acquisition was carried out with MinKNOW (v24.02). Base calling, read quality control, error correction, and adapter trimming were performed using Dorado 0.6.1 on the Genotoul computing cluster (Bioinfo Genotoul, Toulouse Occitanie, France). Genome assembly was performed with Flye (v2.9.5) and gene prediction and annotation with Prokka (v1.14.6), both run on the Genotoul computing cluster. Biosynthetic gene clusters (BGCs) were predicted using antiSMASH v7.1. Core-genome phylogenetic analysis was performed using Panaroo (v1.5) [32] followed by IQ-TREE (v2.3.6) on the core-genome alignment.

### RNA extraction and transcriptomic analysis

Root tissues were ground in liquid nitrogen and total RNA was extracted using the RNeasy Plant Mini Kit (Qiagen) following the manufacturer’s instructions. RNA quality and concentration were verified by NanoDrop (Thermo Fisher Scientific). Library preparation and sequencing were performed using the Illumina TruSeq Stranded mRNA kit and sequenced on an Illumina NovaSeq platform (2 × 150 bp paired-end reads). Read quality was assessed with FastQC (v0.12.1). Reads were aligned with HISAT2 (v2.2.1) to the *Medicago truncatula* reference genome (Mt5.0) and gene expression was quantified as transcripts per million (TPM). Differential expression analysis was performed in R using the DESeq2 package [33]. Differentially expressed genes (DEGs) were defined as |log₂FC| > 1 and FDR < 0.001 in at least one condition, with p-values adjusted using the Benjamini–Hochberg procedure.

### Quantitative RT PCR

cDNA was synthesized from 1 µg total RNA using the High-Capacity cDNA MultiScribe Reverse Transcription Kit (Thermo Fisher Scientific). qPCR was performed on a Bio-Rad CFX using SYBR Green chemistry. *A. euteiches* infection was quantified using tubulin primers (forward: CGGCTCTGGTTTGGGTAGT; reverse: AACCGAGCTTGCTCTTGCG) normalized to *M. truncatula* EF1α (forward: TAACAAGATGGATGCTACC; reverse: GATTTCATCGTACCTAGCCTTTGA). DAPG-producing bacteria were quantified using *phlD*-specific primers designed in this study from a consensus alignment of the 20 top-scoring *phlD* sequences retrieved from NCBI for the genus *Pseudomonas* and aligned with ClustalW (forward: CCTGGGGTТGATAGCAGAGC; reverse: TACAACTGCCCATCGCTCAA; amplicon 141 bp; Tm ∼60°C), normalized to *M. truncatula* EF1α. Cycling conditions were: 95°C 2 min; 40 cycles of 95°C 15 s / 60°C 1 min. Relative expression was calculated using the 2^⁻ΔΔCT^ method.

### Untargeted metabolomics

Metabolites were extracted from the flowpot zeolite substrate by performing five successive washes with 10 mL of 50% methanol containing 0.05% formic acid (Fisher Chemicals, analytical grade). The combined flowthrough was concentrated on a rotary evaporator (Rotavapor R-300, BüCHI) at 40°C and dried under nitrogen flow (N-EVAP 112, Organomation). Dry extracts were resuspended in 100% methanol at 2 mg mL⁻¹ for LC-MS analysis. All extracts were filtered through 0.2 µm PTFE filters (Thermo Scientific) prior to injection. Ultra-high-performance liquid chromatography−high-resolution MS (UHPLC−HRMS) analyses were performed on a Q Exactive Plus quadrupole (Orbitrap) mass spectrometer, equipped with a heated electrospray probe (HESI II) coupled to a U-HPLC Vanquish H (Thermo Fisher Scientific, Hemel Hempstead, U.K.). Reverse phase separation was done on a Luna Omega Polar C18 column (150 mm × 2.1 mm i.d., 1.6 μm, Phenomenex, Sartrouville, France) equipped with a guard column. The mobile phase A (MPA) was water with 0.05% formic acid (FA), and the mobile phase B (MPB) was acetonitrile with 0.05% FA. The solvent gradient was 98% MPA (0 – 0.5 min), 98% MPA to 30% MPB (0.5 - 8 min), 98% MPB (8 - 9 min), 98% MPB (9 - 12 min), 98% MPA (12.1 – 14 min). The flow rate was 0.4 mL/min, the column temperature was set to 40 °C, autosampler temperature was set to 10 °C, and injection volume fixed to 2 μL.

Mass detection was performed in positive and negative ionization (PI and NI) mode at resolution 35 000 power [full width at half-maximum (fwhm) at 400 m/z] for MS1 and 17 500 for MS2 with an automatic gain control (AGC) target of 1 × 10^6^ for full scan MS1 and 1 × 10^5^ for MS2. Ionization spray voltages were set to 3.5 kV for PI and 2,6 kV for NI, and the capillary temperature was kept at 300 °C. The mass scanning range was m/z 100−1500. Each full MS scan was followed by data-dependent acquisition of MS/MS spectra for the four most intense ions using stepped normalized collision energy of 20, 40, and 60 eV.

UHPLC-HRMS raw data were processed using MZmine version 4.7 [34] to extract mass signals between 100 and 1500 Da from 0.5 to 12 min. Respectively, MS1 and MS2 tolerances were set to 10 and 15 ppm in centroid mode. The optimized detection threshold was set to 1 × 106 concerning MS1 and 10 for MS2 using Local minimum feature resolver. Isotope and MS1 scans without MS2 were removed before peak list alignment using join aligner module with a retention time tolerance of 0.1 minute. Ion linkage was deconvoluted using ion identity network using the default adduct and neutral loss list. Mass spectral similarity network was calculated using the modified cosine algorithm with a threshold of 0.7. Spectral library search was used for annotation level 1 using internal AgromiX database with a similarity score of 0.85 and RT tolerance of 0.2. For annotation level 2, the last version of FragHub database [35] was used using a similarity threshold of 0.7. Annotation levels 3 and 4 were performed with Sirius CSI v 6.2 [36] with 10 ppm tolerance in global configuration and spectral matching on FragHub for analog search. Molecular formulas were deciphered using Database search with C,H,N,O,P and S for elemental composition. Properties and structure databases were ticked using bio database and pubchem as fallback.

Output files from MZmine and Sirius CSI were imported into [37]. Among the top 10 in silico candidates per feature, those matching taxonomic criteria (genus, family) were elevated to Level 3a. Features were filtered using RT clustering (ΔRT ≤ 0.01 min). Within each cluster, the top 2 features by network degree and the top 2 by peak intensity were retained. The Mass Spectral Similarity network was constrained to edges with cosine similarity ≥ 0.7 and ΔRT ≤ 8 min between connected nodes. High-confidence annotations (Levels 1, 2a, 3a) seeded the MSS network for iterative annotation propagation.

For each feature pair, candidate structures were ranked using a weighting parameter set to α = 0.3, prioritizing structural-spectral evidence (70%) over in silico ranking (30%). The top 5 candidates by Link Score were retained per feature before looping through the entire MSS network. Redundant annotations with identical InChIKey identifiers and Pearson correlation among samples > 0.7 were consolidated by selecting the candidate with the highest mean peak height. Positive and negative mode feature lists were merged using ΔRT ≤ 0.05 min, Δm/z ≤ 0.002 Da, and a minimum Pearson correlation ≥ 0.6 across sample intensities. Final annotations were enriched with chemical ontology classifications from ClassyFire (kingdom, superclass, class, subclass) and NPClassifier (pathway, superclass, class).

Multivariate statistical analyses (PCA, PLS-DA, sPLS-DA) were performed using MetaboAnalyst 6.0 [38]. All data were peak-intensity normalized by total sum and autoscaled before multivariate and univariate analysis. For multivariate analysis, Principal component analysis (PCA) and Partial Least Squares - Discriminant Analysis (PLS-DA) were performed, while for univariate analysis, one-way Analysis of Variance (ANOVA) accompanied by Fisher’s least significant difference (Fisher’s LSD) post-hoc analysis (p-value < 0.05).

### DAPG quantification by HPLC-UV

DAPG quantification was performed by reversed-phase HPLC coupled to UV detection (Ultimate 3000, Thermo Scientific). Separation was achieved on an XBridge C18 column (25 cm × 4.6 mm × 5 µm, Waters) protected by an XBridge guard column (2 cm × 4.6 mm × 5 µm, Waters). An injection volume of 10 µL was used at a constant flow rate of 0.8 mL min⁻¹. Separation was performed with a gradient of solvent A (water + 0.1% formic acid) and solvent B (acetonitrile + 0.1% formic acid) as follows: 2 min at 20% B; 15.5 min gradient from 20 to 98% B; 9.5 min at 98% B; 2 min gradient from 98 to 20% B; re-equilibration in 7 min. UV detection was set at 270 nm for DAPG and 286 nm for MAPG. Quantification was performed using Chromeleon 7.2 software (Thermo Scientific) based on calibration curves generated from pure DAPG and MAPG standards analyzed under the same conditions.

### Induction of DAPG production by *A. euteiches* culture filtrate

*Aphanomyces euteiches* cell-free culture filtrate was prepared by incubating ten 6 mm mycelium plugs from a 7-day-old CMA culture in 10 mL of PDB in a 50 mL Falcon tube for 3 days at 22°C in darkness, then the culture medium was recovered by centrifugation (4,000 rpm, 10 min) and filtered through a 0.22 µm membrane. DAPG-producing strain A05H was pre-cultured overnight in PDB at 28°C with shaking (200 rpm) and adjusted to OD_600_ = 0.1 and was then inoculated into fresh PDB supplemented with increasing proportions of *A. euteiches* culture filtrate (0, 25, 50, 100% v/v) and grown for 24 h at 28°C (n = 8–16 replicates per condition). At the end of the incubation, the bacteria culture medium was recovered by centrifugation and 0.22 µm filtration, and DAPG was quantified directly in the crude filtered medium by HPLC-UV as described above, with concentrations normalized to the injected volume of medium. the *A. euteiches* filtrate was subjected to heat treatment (95°C, 10 min) and size-based fractionation by successive ultrafiltration through 100 kDa and 30 kDa molecular weight cut-off membranes (Amicon, Merck). Statistical differences were assessed by one-way ANOVA followed by Tukey’s HSD post-hoc test.

### Statistical Analysis

All statistical analyses were performed in R (v4.6.0). One-way ANOVA followed by Tukey’s HSD post-hoc test was used for comparisons between more than two groups. Unpaired two-tailed Student’s t-tests were used for pairwise comparisons. Community-level differences were assessed by PERMANOVA (999 permutations) on Bray-Curtis distance matrices using the vegan package. Differential abundance between conditions was performed with DESeq2 using the Wald test with Benjamini-Hochberg adjustment. A significance threshold of α = 0.05 was applied throughout.

## Results

### Construction of a functionally diverse rhizosphere SynCom from *M. truncatula*

To construct a representative synthetic community (SynCom) of *M. truncatula* F83000.5, 1,364 bacterial strains were isolated from the rhizosphere of plants cultivated in pot soil using a limiting dilution approach (Fig.1A). High-throughput 16S rRNA gene sequencing on an Illumina MiSeq platform generated 1,076,049 high-quality reads across the 1,364 strains. Reads were clustered into operational taxonomic units (OTUs) at 97% sequence similarity using QIIME2 with VSEARCH, yielding 355 OTUs. After quality filtering and retaining only wells containing a single OTU, 812 wells corresponding to 79 distinct OTUs (clustered at 97% sequence similarity) were retained for downstream analysis (Fig. 1A; Supplementary Table S1).

**Figure 1.**
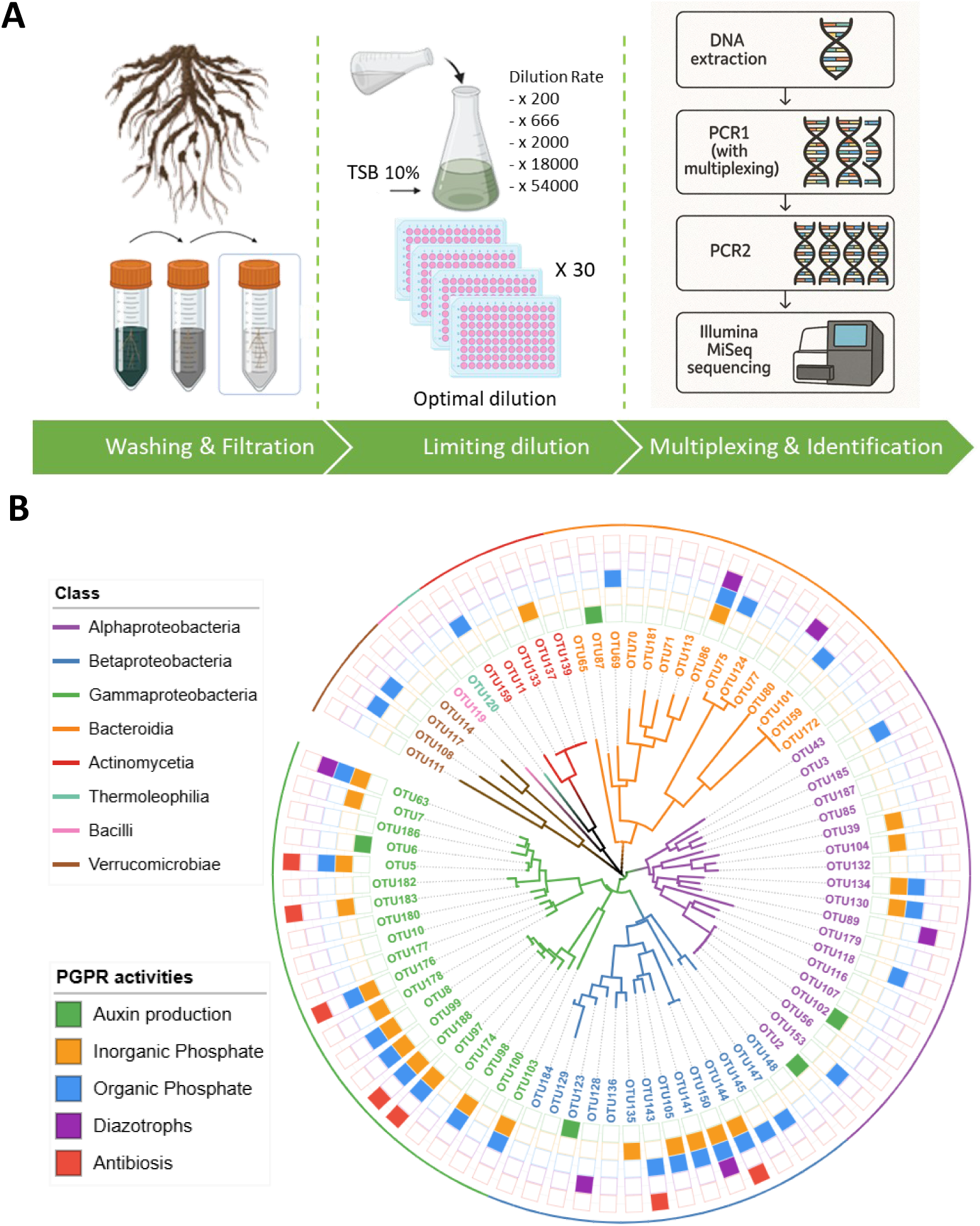
Isolation pipeline and functional characterization of the *M. truncatula* rhizosphere SynCom. **A.** Schematic overview of the bacterial isolation workflow. Root samples from *M. truncatula* were subjected to washing and filtration to recover rhizosphere-associated bacterial cells. The resulting suspension was serially diluted in TSB and distributed into microtiter plates following a limiting dilution strategy to favor single-cell isolation. Bacterial DNA was extracted from positive wells, amplified by two successive PCR steps and sequenced on an Illumina MiSeq platform. **B.** Phylogenetic tree of the OTUs identified within the cultivable rhizosphere community. Branch colors indicate bacterial class affiliation (see legend). Colored squares on the outer ring indicate plant growth-promoting rhizobacteria (PGPR) activities detected among isolates selected for the SynCom: auxin production (green), inorganic phosphate solubilization (orange), organic phosphate (phytate) solubilization (blue), diazotrophic potential (purple), and antagonistic activity against *A. euteiches* (red). OTUs bearing no colored squares were included in the SynCom based on taxonomic representativeness alone.

Phylogenetic analysis of the 79 strains revealed a broad taxonomic diversity spanning eight bacterial classes, with Gammaproteobacteria, Betaproteobacteria, and Alphaproteobacteria as the dominant lineages, alongside representatives of Bacteroidia, Actinomycetia, Thermoleophilia, Bacilli, and Verrucomicrobiae (Fig. 1B). The three most abundant genera were *Rhodanobacter*, the most abundant genus accounting for 384,061 reads across 519 occurrences, followed by *Pseudomonas* (299,716 reads, 416 occurrences) and *Dokdonella* (136,592 reads, 226 occurrences).

From this collection, a SynCom was assembled by selecting isolates prioritizing both taxonomic representativeness and functional complementarity, with a particular emphasis on activities relevant to plant protection and nutrition. The selected strains collectively cover the full taxonomic breadth of the cultivable community and display a diverse array of plant-beneficial traits (Fig. 1B; Supplementary Table S1). These include phosphate solubilization activity — both inorganic (orange squares) and organic phytate (blue squares) — auxin production (green squares), diazotrophic potential (purple squares), and direct antagonistic activity against *A. euteiches* (Fig. 1B; red squares). Notably, strains with anti-*Aphanomyces* activity were rare in the overall collection but were systematically included in the SynCom given their potential relevance to disease suppression.

### *Aphanomyces euteiches* infection selectively enriches *Pseudomonas* strains in the rhizosphere

To determine the impact of *A. euteiches* infection on the SynCom composition, plants were cultivated in a flow pot system and inoculated with the SynCom followed by inoculation with *A. euteiches* zoospores. Full-length 16S rRNA amplicon sequencing (Oxford Nanopore Technology, NanoClust pipeline) was performed on rhizosphere samples collected 21 days post-inoculation. Principal Coordinates Analysis (PCoA) based on Bray-Curtis dissimilarity revealed a clear separation of the three conditions along Axis 1, which explained 81.3% of the total variance (Fig. 2A). Community composition differed significantly across conditions (PERMANOVA, p = 0.001, ***). Alpha diversity analyses further revealed a progressive reduction in both diversity and richness across conditions (Fig. 2B). Shannon diversity index decreased significantly from the SynC_T0 inoculum (∼3.0) upon plant colonization (Mt_SynC, ∼2.2) and even more markedly under infection (Mt_Ae_SynC, ∼1.7; Tukey groups a, b, c). A similar trend was observed for observed richness (∼45, ∼34, and ∼30 OTUs respectively; groups a, b, b). Taxonomic profiling revealed that this reduction in diversity was associated with a dramatic compositional shift: *A. euteiches* infection triggered a strong enrichment of *Pseudomonas*, which became the dominant taxon at the expense of other genera including *Rhodanobacter*, *Variovorax*, and *Methylobacillus* (Fig. 2C; Supplementary Table S2).

**Figure 2.**
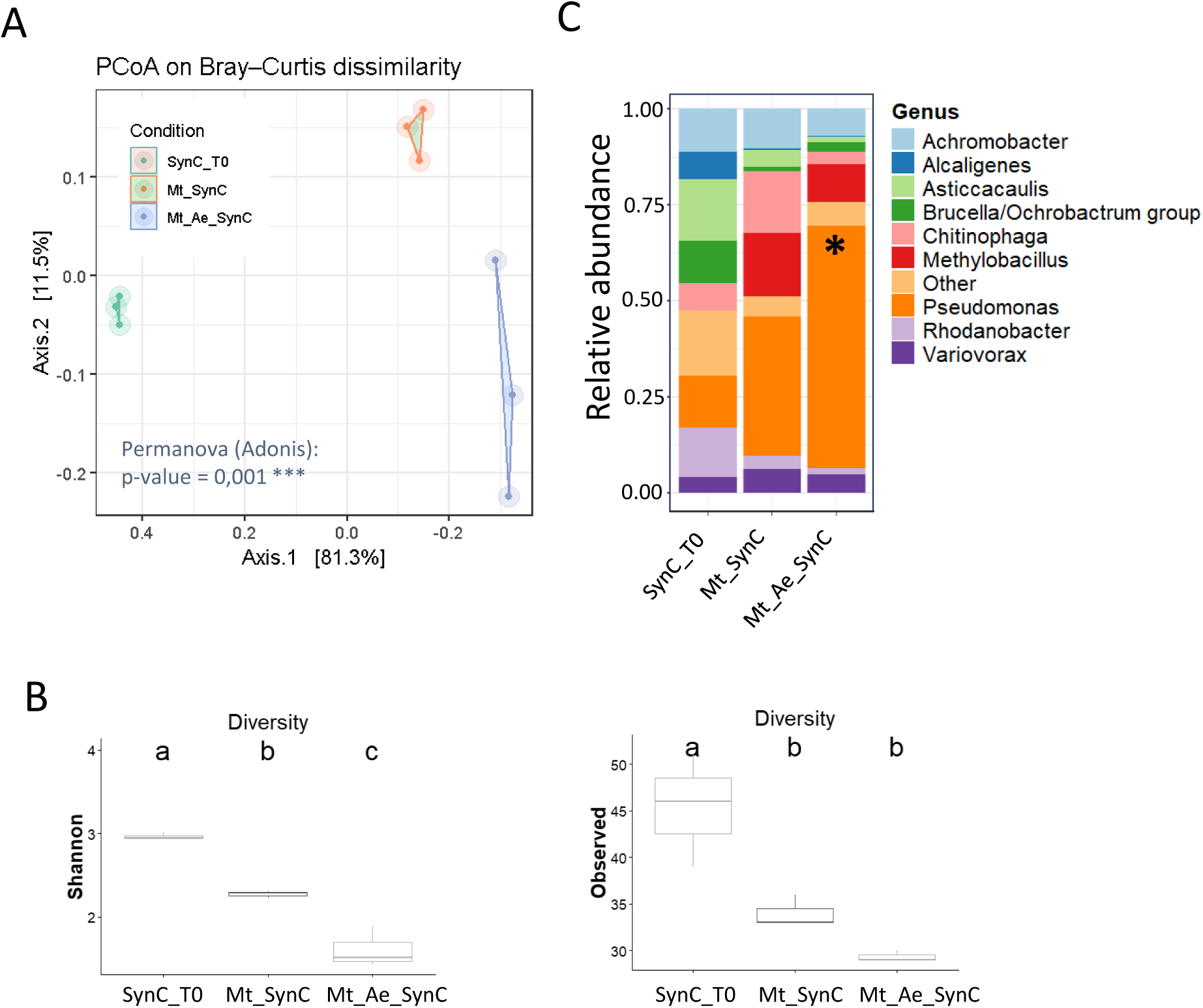
Impact of *A. euteiches* infection on the SynCom structuration. Full-length 16S rRNA amplicon sequencing was performed on rhizosphere samples collected 21 days post-inoculation from three conditions: SynC_T0 (inoculum prior to plant contact), Mt_SynC (*M. truncatula* colonized by the SynCom without pathogen), and Mt_Ae_SynC (*M. truncatula* colonized by the SynCom and infected with *A. euteiches*). **A.** Principal Coordinates Analysis (PCoA) based on Bray-Curtis dissimilarity. Each point represents one biological replicate (n = 3 per condition). Community composition differed significantly across conditions (PERMANOVA/Adonis, p = 0.001, ***). **B**. Alpha diversity metrics across conditions. Left panel: Shannon diversity index. Right panel: observed richness (number of OTUs). Different letters (a, b, c) indicate statistically significant differences between groups (p < 0.05). C. Stacked bar plots showing the relative abundance of the dominant bacterial genera across conditions. Each bar represents the mean relative abundance of three biological replicates. Asterisk (*) indicates a significant enrichment of *Pseudomonas* in the Mt_Ae_SynC condition relative to Mt_SynC.

Collectively, these data showed that *A. euteiches* infection reduced in community diversity and increase the abundance of *Pseudomonas* strains, consistent with a disease-driven recruitment of potentially antagonistic bacteria.

### *Aphanomyces* infection induces a dual defense response involving both plant- and microbiota-derived metabolites

To determine whether the pathogen-induced restructuring of the SynCom translated into functional metabolic changes, we performed untargeted metabolomic profiling of *M. truncatula* rhizosphere across four conditions: plant alone (Mt), plant infected with *A. euteiches* (Mt_Ae), plant colonized by the SynCom (Mt_SynC), and the tripartite interaction (Mt_Ae_SynC). A total of 595 variables were retrieved across all the samples (Supplementary Table S3). Partial Least Squares Discriminant Analysis (PLS-DA) revealed a clear separation of all four conditions (Fig. 3A). The primary separation was driven by pathogen infection (44%), whereas SynCom colonization further discriminated infected from non-infected samples (20%), resulting in a distinct metabolic profile for the tripartite interaction demonstrating that the SynCom significantly reshapes the metabolic landscape of infected roots beyond the plant response alone.

**Figure 3.**
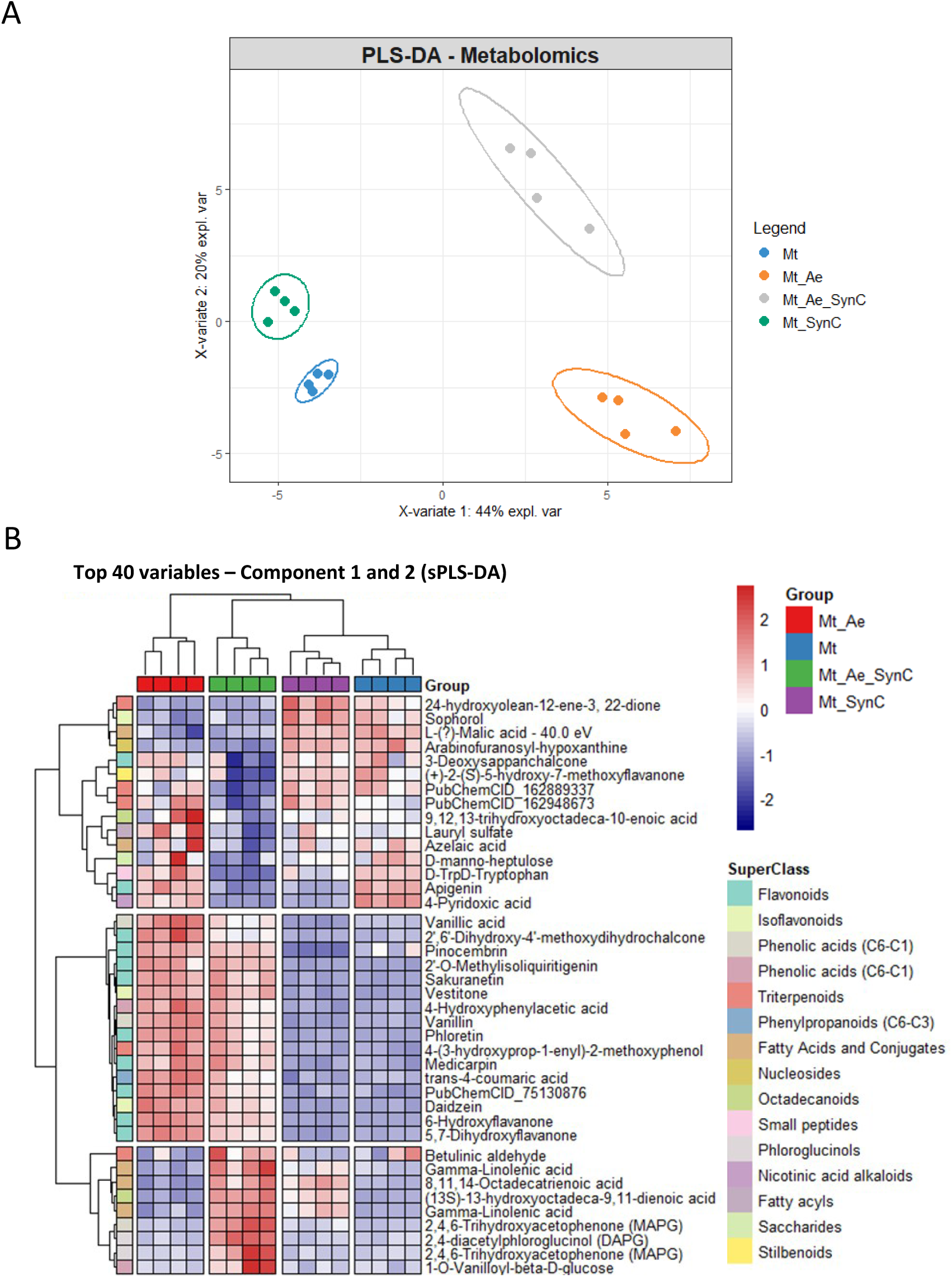
Untargeted metabolomic profiling reveals a dual defense response in *A. euteiches*-infected *M. truncatula* roots colonized by the SynCom. **A.** Partial Least Squares Discriminant Analysis (PLS-DA) of root metabolomes across four conditions: Mt (plant alone, dark blue), Mt_Ae (plant infected with *A. euteiches*, orange), Mt_SynC (plant colonized by the SynCom, green), and Mt_Ae_SynC (tripartite interaction, grey). X-variate 1 (44% of explained variance) separates conditions primarily by infection status; X-variate 2 (20%) captures the additional effect of the SynCom. Each point represents one biological replicate. Ellipses represent 95% confidence intervals. **B.** Heatmap of the top 40 discriminant metabolites identified by sparse PLS-DA (sPLS-DA) on components 1 and 2. Rows represent individual metabolites annotated by chemical superclass (right color bar). Columns represent biological replicates grouped by condition. Color scale reflects scaled metabolite abundance (red: high; blue: low). Two major clusters are distinguishable: an isoflavonoid/phenylpropanoid cluster enriched in infected conditions (Mt_Ae and Mt_Ae_SynC), and a phloroglucinol cluster — including DAPG and its precursor MAPG — exclusively enriched in the tripartite condition (Mt_Ae_SynC).

Sparse PLS-DA (sPLS-DA) hierarchical clustering of the top 20 discriminant variables resolved two major metabolic modules associated with disease condition (Fig. 3B). The first module, enriched in the Mt_Ae condition, comprised a suite of isoflavonoid and phenylpropanoid phytoalexins characteristic of legume immune activation [39], including the major phytoalexin medicarpin, its precursor vestitone, upstream isoflavones such as daidzein and 6-hydroxyflavanone, chalcone-derived compounds including sakuranetin, pinocembrin, phloretin and 2-O-methylisoliquiritigenin, and phenolic acids such as trans-p-coumaric acid and vanillic acid. Importantly, this module remained strongly activated in the tripartite condition, demonstrating that the SynCom does not suppress the plant’s endogenous chemical defenses. The second module was specifically and exclusively enriched in the tripartite condition Mt_Ae_SynC. It contained three metabolites related to the oxylipin pathway (linolenic acid, 13-hydroxyoctadeca-9,11-dienoic acid, and octadecatrienoic acid), a pathway which was suggested to play a role in the defense response against *A. euteiches* [16, 40]. This cluster also includes a triterpenoid, the betulinic aldehyde. Triterpenoids are produced by legume plants in large quantity and they could play a role in the interaction with microorganisms [41, 42]. Finally, we observed also the production of the broad-spectrum antibiotic 2,4-diacetylphloroglucinol (DAPG), a metabolite produced *Pseudomonas* species and its biosynthetic precursor monoacetylphloroglucinol (MAPG, annotated as 2,4,6-trihydroxyacetophenone). The selective accumulation of DAPG and MAPG exclusively in infected roots colonized by the SynCom, but not in healthy SynCom-colonized roots (Mt_SynC), suggests that DAPG biosynthesis is conditionally activated during plant infection. Taken together, these data identified two distinct metabolomic signature associated with *A. euteiches* infection: a direct plant defense response to *A. euteiches* then potentiated by a pathogen-triggered bacterial contribution to plant defense through DAPG production.

### The rhizosphere SynCom amplifies plant immune responses to *A. euteiches*

To determine the impact of the SynCom on the transcriptional response of *M. truncatula* roots *A. euteiches* infection, we profiled root transcriptomes across four conditions: mock plants, SynCom alone (SynC), *A. euteiches* alone (Ae), and the tripartite interaction (Ae_SynC). Differential gene expression analysis detected 5,773 DEGs (Log2 FC <-1 or >1 and FDR <0.001 in at least one condition; Supplementary Table S4). In more details 221, 4321 and 4596 DEGs specific to SynC, Ae and Ae_Sync respectively were detected, revealing distinct transcriptional signatures depending on the condition (Fig. 4A). Colonization of the host plant by the SynCom alone triggered a limited response, with very few DEG across all Log2 fold-change classes (48, 58, 24, and 3 genes for classes Log2 fold change >1–<2, >2–<3, >3–<4, and >4, respectively), demonstrating that the SynCom establishes stable root colonization without imposing a significant transcriptional response on the host. In contrast, infection with *A. euteiches* alone induced a substantially stronger response (504, 364, 246, and 138 genes per class), consistent with the activation of a broad defense program as already reported [39]. Strikingly, the tripartite condition generated the most extensive transcriptional reprogramming with the highest number of differentially expressed genes across every fold-change class (828, 565, 337, and 272 genes respectively), indicating that the simultaneous presence of the SynCom and the pathogen potentiates host responses.

**Figure 4.**
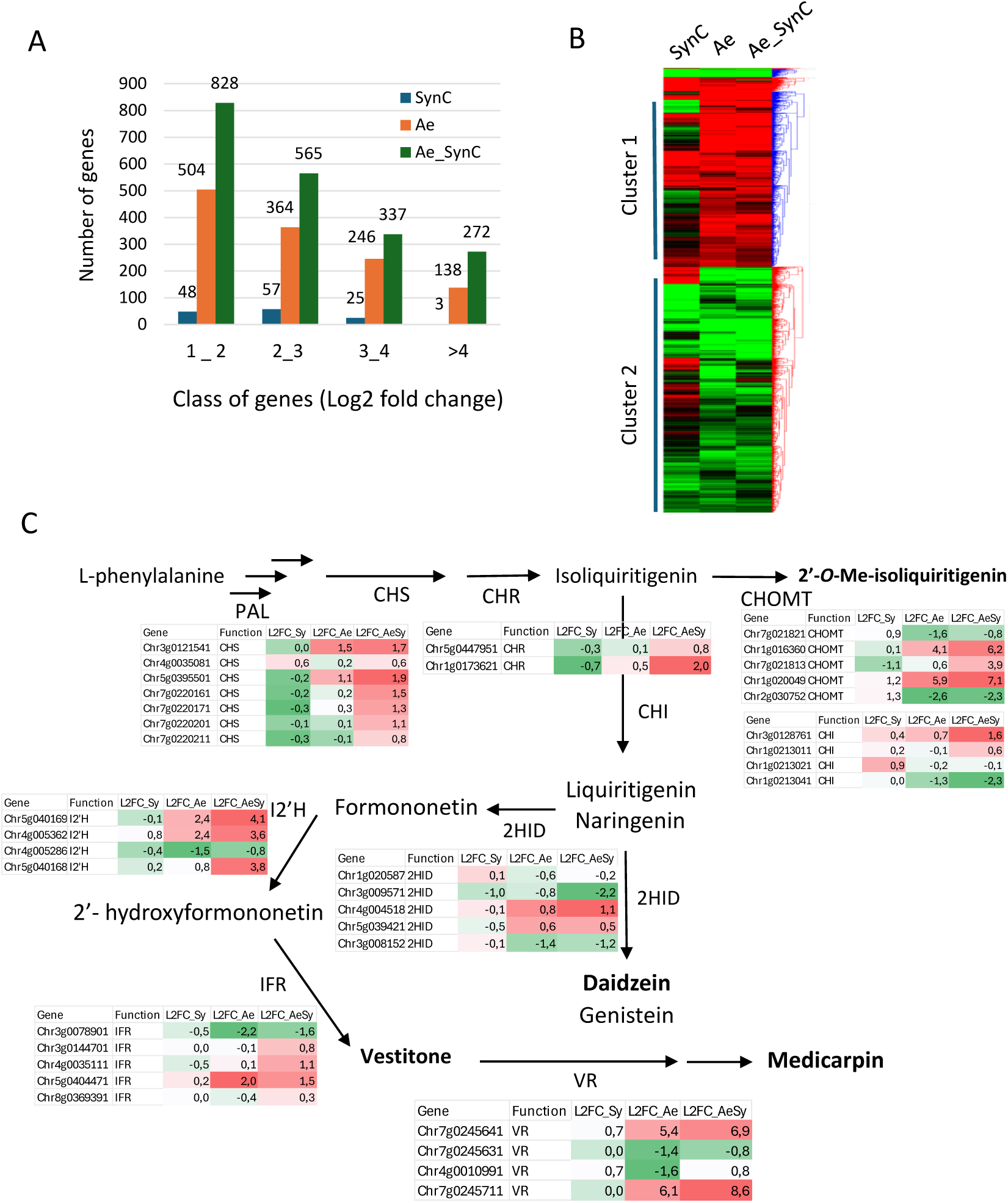
The SynCom potentiates the transcriptional activation of the isoflavonoid defense pathway in *A. euteiches*-infected *M. truncatula* roots. **A.** Number of differentially expressed Number of differentially expressed genes (DEGs) per Log2 fold-change class in three conditions relative to uninfected, uninoculated controls: SynCom alone (SynC, blue), *A. euteiches* alone (Ae, orange), and the tripartite interaction (Ae_SynC, green). Classes correspond to Log2FC intervals: 1–2, 2–3, 3–4, and >4. **B.** Heatmap of genes differentially expressed across SynC, Ae, and Ae_SynC conditions. Two main clusters are identified: Cluster 1 (genes induced by infection) and Cluster 2 (genes specifically amplified in the tripartite condition). Clustering was performed using the Hierarchical Clustering Explorer program V. 3.5, with normalization by standardization (Mean and Standard Deviation), Average Distance (UPGMA) and Similarity/Distance measure by Euclidean distance. **C.** Schematic representation of the isoflavonoid biosynthetic pathway leading to the phytoalexin medicarpin. Tables at each enzymatic step show the Log2FC of the corresponding gene(s) in SynCom (blue), Ae (orange), and Ae_SynC (green) conditions. Enzymes shown: PAL, phenylalanine ammonia-lyase; CHS, chalcone synthase; CHR, chalcone reductase; CHOMT, caffeic acid O-methyltransferase; CHI, chalcone isomerase; I2’H, isoflavone 2’-hydroxylase; IFR, isoflavone reductase; VR, vestitone reductase. Compounds detected in higher amounts in the rhizosphere upon *A. euteiches* infection are in bold letters.

Hierarchical clustering performed on DEGs led to the definition of two main clusters (Fig. 4B) gathering genes induced during *A. euteiches* infection (Cluster 1; 2283 genes) or repressed after *A. euteiches* infection (Cluster 2; 3207 genes). Mining a selection of the most induced genes belonging to the cluster 1 (Log2 FC>4) revealed the overrepresentation of functional classes related to plant defenses such as transcription factor, defense proteins and secondary metabolism notably belonging to the biochemical pathway leading to the synthesis of isoflavonoids such as the phytoalexin medicarpin (Supplementary table S5). This analysis also revealed that the SynCom colonization alone left these genes essentially silent, with Log2FC values close to zero.

Since the induction of the isoflavonoid pathway is one of the major response of legumes to pathogens, we examined this pathway in more details. Mapping these expression data onto the isoflavonoid biosynthetic pathway illustrated a coordinated, stepwise transcriptional activation specifically reinforced by the tripartite interaction (Fig. 4C; Supplementary Table S6). Entry into the pathway through phenylalanine ammonia-lyase was followed by robust induction of chalcone synthase (CHS) and chalcone reductase (CHR), which direct carbon flux toward the isoflavonoid branch. Downstream, chalcone isomerase (CHI), caffeic acid O-methyltransferase (CHOMT), isoflavone 2′-hydroxylase (I2′H), isoflavone reductase (IFR), and vestitone reductase (VR) — the terminal enzyme leading to the phytoalexin medicarpin — were all significantly more induced in the tripartite condition than in infection alone. For example, VR showed Log2FC values of 0.7, 5.4, and 6.9 in SynCom, Ae, and Ae_SynCom respectively, and a second VR isoform reached L2FC 0.0, 6.1, and 8.6. Collectively, these findings demonstrate that SynCom colonization potentiates the host isoflavonoid biosynthetic pathway in response to *A. euteiches*. This systematic amplification across every step of the pathway demonstrates that the SynCom does not interfere with host immunity but rather enhances the plant’s capacity to produce medicarpin and upstream isoflavone intermediates in response to pathogen challenge.

### *Aphanomyces euteiches* infection selectively enriches antagonistic DAPG-producing *Pseudomonas*

Metabarcoding analysis of the SynCom during *A. euteiches* revealed an increase of abundance of *Pseudomonas* strains. The SynCom contained seven OTUs associated to the *Pseudomonas* genus (A05H, D08H, E12A, F01D, G10J, A12A and A08K), a genus repeatedly associated with disease-suppressive activity in the rhizosphere and DAPG production. To identify the strains able to produce DAPG, the presence of the *phlD* gene encoding the key enzyme of the DAPG biosynthetic pathway was screened in the seven strains by PCR. Only A05H and D08H carried *phlD*, identifying them as the only potential DAPG producers of the community. To characterize the *Pseudomonas* strains at the genomic level, the genomes of A05H and D08H were sequenced resulting in the assembly of a single contig corresponding to a complete genome of 6.67 Mb highly similar between the two strains. Prediction of Biosynthetic Gene Cluster (BGC) using antiSMASH confirmed the presence of a complete *phlD* locus in the genome of A05H and D08H, together with BGCs for hydrogen cyanide (HCN) and the siderophore histicorrugatin (Supplementary Table S7). A core-genome phylogenetic analysis placed A05H and D08H in a clade closely related to *P. brassicacearum* LMG 21623 and *P. ogarae* F113, two well-characterized DAPG producing strains (Fig. 5A).

**Figure 5.**
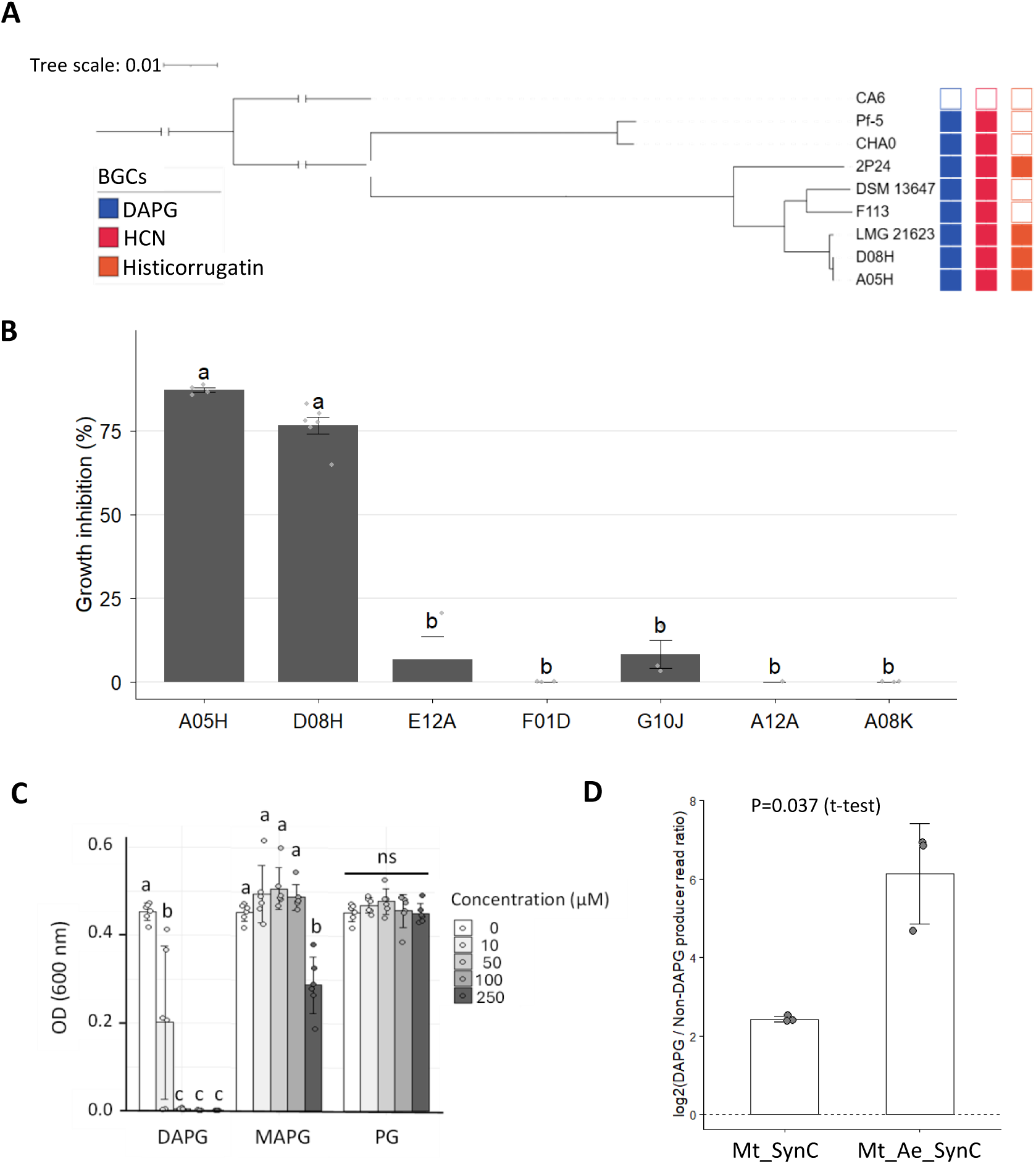
Genomic and biological characterization of DAPG-producing *Pseudomonas* strains from the SynCom. **A.** Core-genome phylogenetic tree of SynCom DAPG producing *s* strains A05H and D08H and related reference strains, reconstructed from a Panaroo-based core-genome alignment. Scale bar = 0.01 substitutions per site. Bootstrap values (%) are shown at nodes. Colored squares indicate predicted biosynthetic gene clusters (BGCs): DAPG (blue), HCN (red), histicorrugatin (orange). **B.** Antagonistic activity of SynCom *Pseudomonas* strains against *A. euteiches* in dual co-culture assays on PDA medium. Bars represent percentage of mycelial growth inhibition relative to the no-bacteria control. Different letters indicate statistically significant differences (ANOVA, Tukey post-hoc, p < 0.05). **C.** Dose-dependent inhibition of *A. euteiches* growth by purified DAPG, measured by optical density (OD). Increasing DAPG concentrations (0, 10, 50, 100, 200 µM) were applied to *A. euteiches* cultures. Different letters denote significant differences between concentrations (ANOVA, Tukey, p < 0.05). **D.** Relative enrichment of DAPG-producing strains, expressed as log₂(DAPG-producer/non-DAPG-producer read ratio) derived from 16S metabarcoding reads mapped to SynCom member genomes, in the absence (Mt_SynC) or presence of *A. euteiches* (Mt_Ae_SynC). The ratio is significantly higher in the infected condition (t-test, p = 0.037), indicating pathogen-driven enrichment of DAPG producers. Individual points represent biological replicates; bars show means ± SD.

When the ability of the seven *Pseudomonas* strains to inhibit *A. euteiches* was assessed in dual confrontation assays on solid media, only A05H and D08H strongly inhibited mycelial growth, whereas the five *phlD*-negative *Pseudomonas* strains showed no significant activity (Fig. 5B). Purified DAPG, but not the DAPG precursors MAPG and PG likewise inhibited *A. euteiches* growth in a dose-dependent manner (Fig. 5C), directly linking the *phlD* genotype to the antagonistic phenotype.

Since the HCN (volatile compound) and histicorrugatin (siderophore) BGCs were present in the two strains displaying anti- *A. euteiches* activity, we assessed volatile-mediated inhibition using a split-plate system preventing direct contact, and iron-dependent inhibition by supplementing the medium with FeCl_3_ to saturate siderophore activity. In the absence of direct contact, *Aphanomyces* hyphae grow normally, indicating that bacterial volatile compounds are not involved in growth inhibition and the growth of *A. euteiches* remains strongly impacted despite the presence of iron in the medium, indicating that siderophore activity is not involved in the antagonistic activity of A05H and D08H. These results indicate that DAPG is the primary bioactive compound mediating the antagonistic activity of A05H and D08H (Supplementary Fig. S1). Finally, to determine whether DAPG-producing strains were selectively enriched in planta, metabarcoding reads were mapped onto the *Pseudomonas* genomes and computed the ratio of DAPG-producer to non-DAPG-producer reads. This ratio was significantly higher under *A. euteiches* infection (Mt_Ae_SynC) than in the uninfected condition (Mt_SynC) (Fig. 5D; t-test, p = 0.037), demonstrating that infection selectively increased the abundance of the DAPG-producing strains within the *Pseudomonas* population.

### DAPG-producing *Pseudomonas* strains reduce disease symptoms

To directly assess the contribution of DAPG-producing *Pseudomonas* strains to disease suppression, we conducted a pot experiment using vermiculite substrate under conditions comparable to the flowpot system. *Medicago truncatula* plants were inoculated with either a complete SynCom containing the two DAPG-producing strains D08H and A05H (SynCphl) or a SynCom depleted of these strains (SynC-), in the presence or absence of *A. euteiches*. Disease symptoms were evaluated by root phenotyping and pathogen load was quantified by qPCR. Visual inspection of root systems revealed that *A. euteiches* infection caused severe root browning and maceration in the Mt_Ae condition. Similar symptoms were largely maintained in plants inoculated with the DAPG-depleted community (MtAe_SynC-) (Fig. 6A). In contrast, plants colonized by the complete DAPG-producing SynCom (MtAe_SynCphl) displayed markedly healthier root systems, with reduced necrosis and better-preserved root architecture. Quantification of root length confirmed these observations (Fig. 6B). Infection with *A. euteiches* alone significantly reduced root length compared to non-infected control plants (Mt vs MtAe, p < 0.05). Plants inoculated with the complete SynCphl community and infected with the pathogen (MtAe_SynCphl) showed root lengths comparable to healthy controls (ns vs Mt), demonstrating a near-complete restoration of root growth. In contrast, the DAPG-depleted SynCom (MtAe_SynC-) failed to provide equivalent protection, with root lengths remaining significantly reduced relative to MtAe_SynCphl (Fig. 6B).

**Figure 6.**
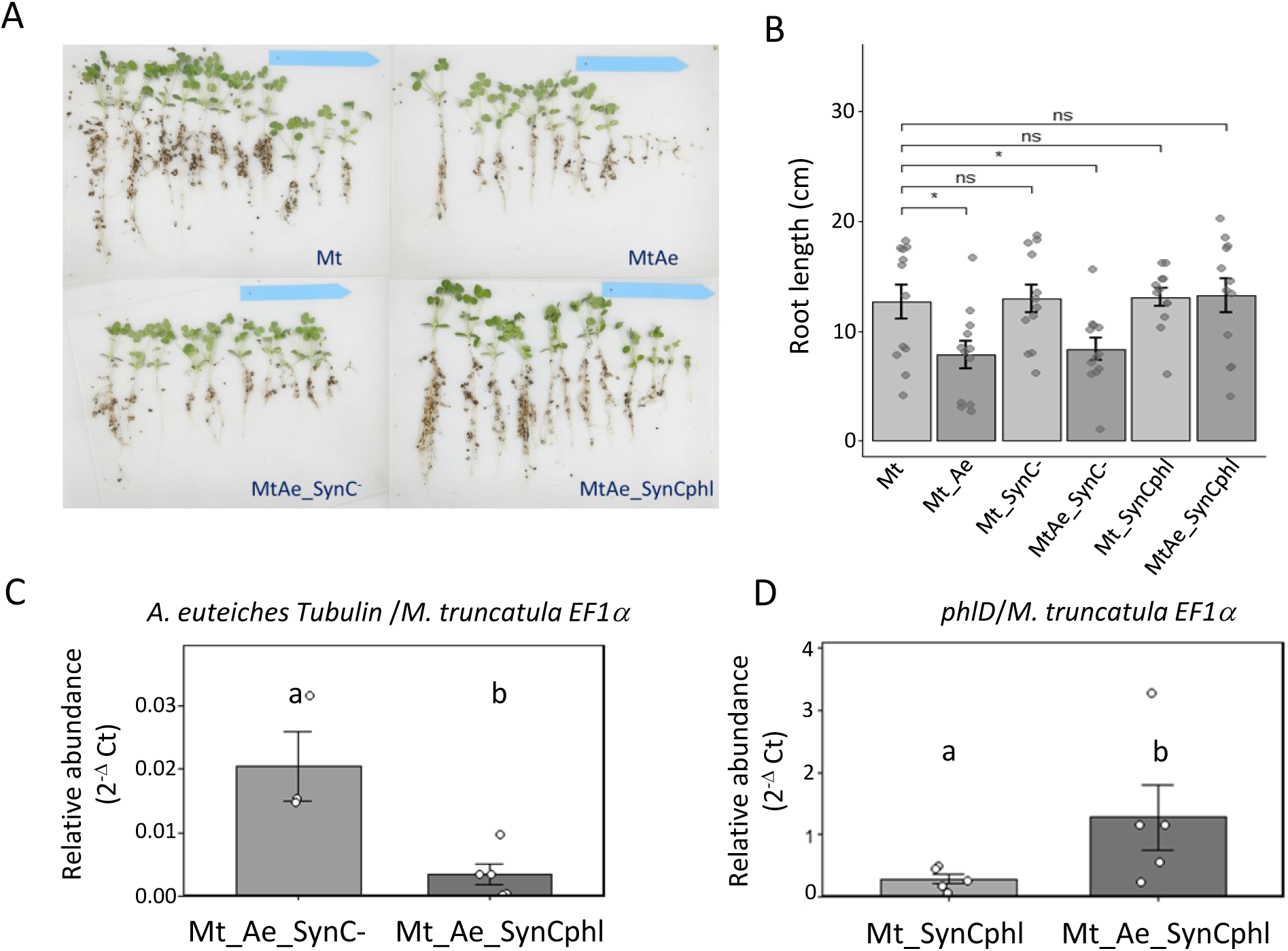
DAPG-producing *Pseudomonas* strains reduce *A. euteiches* root colonization and alleviate root rot symptoms in *M.truncatula*. **A.** Representative photographs of *M. truncatula* root systems 21 days post-inoculation across four conditions: Mt (plant alone), MtAe (plant infected with *A. euteiches*), MtAe_SynC- (plant infected with *A. euteiches* and inoculated with a SynCom depleted of DAPG-producing strains), and MtAe_SynCphl (plant infected with *A. euteiches* and inoculated with the complete DAPG-producing SynCom). **B.** Root length (cm) measured across six conditions. Significant differences are indicated by brackets (* p < 0.05; ns: not significant; ANOVA with post-hoc Tukey test). Each point represents one biological replicate. Error bars show standard error of the mean. **C.** Relative abundance of *A. euteiches* in inoculated roots in presence of the SynCom depleted of DAPG-producing strains (Mt_Ae_SynC-) or with the SynCom containing the two DAPG producting strains (Mt_Ae_SynCphl), quantified by qPCR using *A. euteiches* tubulin primers normalized to *M. truncatula* EF1α (2^−ΔCt). Different letters indicate statistically significant differences (p < 0.05). **D.** Relative abundance of DAPG-producing *Pseudomonas* in roots, quantified by qPCR using *phlD*-specific primers normalized to *M. truncatula* EF1α (2^−ΔCt). Different letters indicate statistically significant differences between Mt_SynCphl and MtAe_SynCphl (p< 0.05). Each point represents one biological replicate; error bars show standard error of the mean.

To determine whether the protective effect of SynCphl was associated with reduced pathogen colonization, we quantified *A. euteiches* biomass in roots by qPCR using *A. euteiches* tubulin normalized to *M. truncatula* EF1α (Fig. 6C). Pathogen abundance was significantly lower in MtAe_SynCphl roots compared to MtAe_SynC-roots (groups b *vs* a, respectively), directly linking the presence of DAPG-producing strains with a lower *A. euteiches* colonization. Finally, to confirm that DAPG-producing *Pseudomonas* strains are indeed enriched *in planta* upon infection, we quantified the abundance of *phlD*-producing bacteria using *phlD*-specific qPCR primers normalized to *M. truncatula* EF1α (Fig. 6D). The relative abundance of *phlD*-producing bacteria was significantly higher in MtAe_SynCphl infected roots compared to non-infected Mt_SynCphl controls (groups b vs a), demonstrating that *A. euteiches* infection actively promotes the *in-planta* proliferation of DAPG-producing *Pseudomonas* strains. Taken together, these results established that DAPG-producing *Pseudomonas* strains are key contributors to microbiota-mediated suppression of *A. euteiches* root rot.

### Extracellular compounds produced by *A. euteiches* induce production of DAPG

Since metabolomics analyses of flowpots experiments revealed the production of DAPG only in pots inoculated with *A. euteiches*, we next sought to determine whether *A. euteiches* directly modulates DAPG production. To determine whether DAPG production requires pathogen-derived signals, we used the experimental setup describes on the figure 7A. The inducing activity on the inhibitory activity against *A. euteiches* was first tested using known inducers of DAPG production, the precursors PG and MAPG [43] which did not have inhibitory activity toward *A. euteiches* (Fig. 5C). A05H culture supernatant not supplemented with MAPG or PG failed to inhibit *A. euteiches* growth confirming that the anti-Aphanomyces activity is inducible (Fig. 7B) whereas cultivation of A05H cells in presence of PG or MAPG induced a strong anti-Aphanomyces activity. Then, we investigated whether compounds released by *A. euteiches* could stimulate the anti-Aphanomyces activity and DAPG production. Mycelium of *A. euteiches* was cultivated in PDB and after 4 days of incubation, extracellular culture medium was added to a suspension of A05H cells. Addition of increasing proportions of *A. euteiches* culture medium (0% to 100%) to A05H cultures triggered a strong, dose-dependent induction of an inhibitory activity toward *A. euteiches* and induced DAPG production (Fig. 7C and D; ANOVA, p = 2.2×10⁻¹³). DAPG levels remained low at 0% and 25% filtrate but increased significantly at 50% and peaked at 100% filtrate, with significant pairwise differences confirmed by post-hoc testing (Fig. 7D). In parallel, the same concentration gradient caused a significant dose-dependent inhibition of *A. euteiches* growth (Fig. 7C; Tukey groups a, b, c, d).

**Figure 7.**
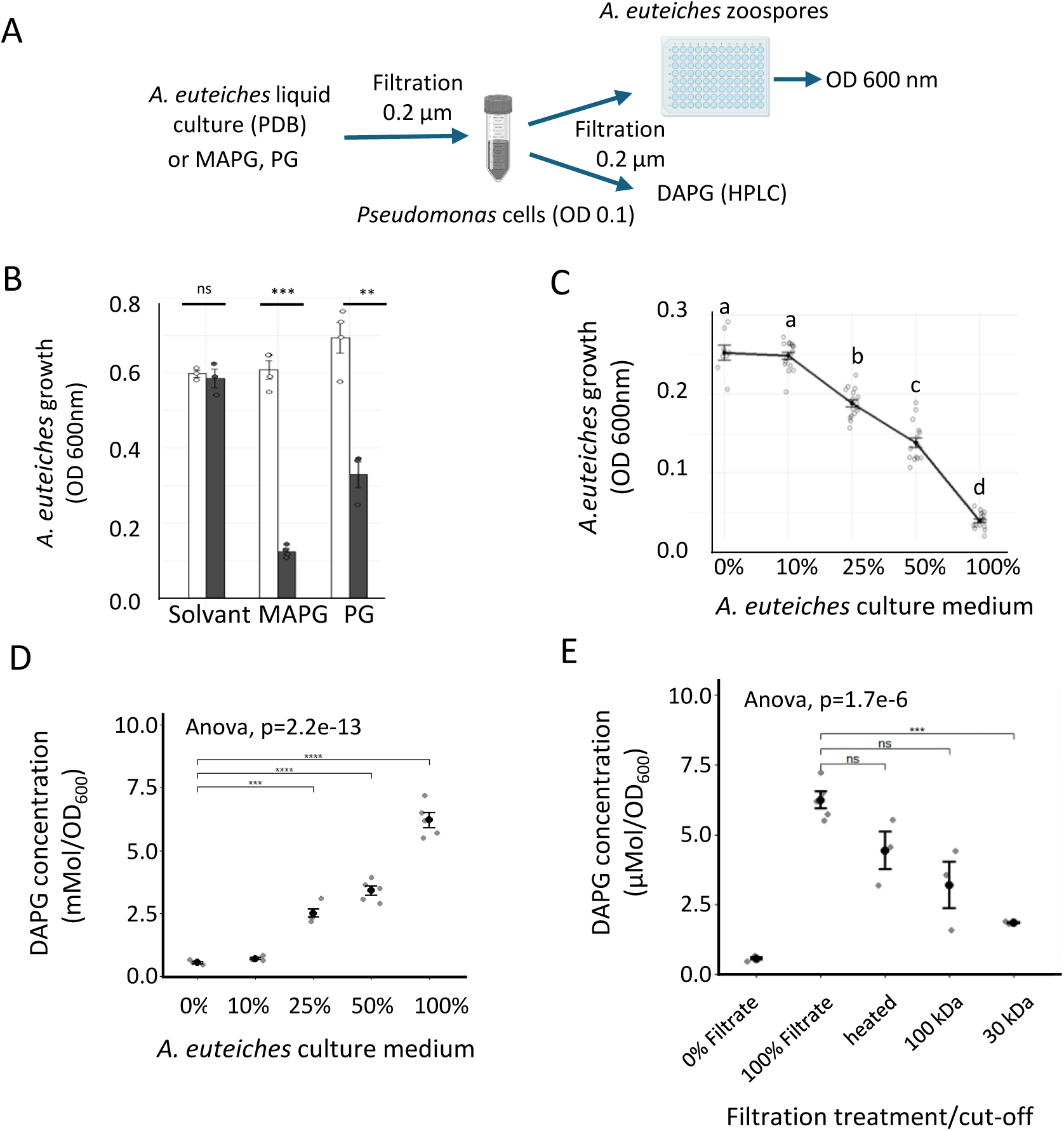
Extracellular compounds released by *Aphanomyces euteiches* induce DAPG production in *Pseudomonas* strain A05H. **A**. Experimental setup. A05H cells were incubated with known inducers of DAPG synthesis (MAPG, PG) or *A. euteiches* culture medium. This medium was first filtered using a 0.22 µm filter to remove mycelial debris, and eventually fractionated using filters with a cut-off of 30 kDa or 100 kDa and tested on growth of *A. euteiches* zoospores and DAPG concentration was determined. **B.** Induction of inhibitory activity toward *A. euteiches* growth in the A05H strain cultivated in the presence of DAPG precursors MAPG or PG (black bars). The white bars correspond to the medium without bacteria. Error bars represent standard error of the mean. **C.** Induction of inhibitory activity toward *A. euteiches* growth in the A05H strain cultivated in the presence of increasing proportions (0–100%) of cell-free *A. euteiches* culture medium. Different letters indicate statistically significant differences between concentrations (ANOVA, post-hoc Tukey, p < 0.05). Each point represents an independent biological replicate. **D.** Normalized DAPG production (µmol/OD_600_) by A05H strain exposed to increasing proportions of *A. euteiches* culture medium. DAPG was quantified by LC-UV. Brackets indicate significant pairwise comparisons relative to the 0% control (*** p < 0.001, **** p < 0.0001; ANOVA, p = 2.2×10⁻¹³). **E.** Normalized DAPG production (µmol/OD_600_) by A05H cells exposed to *A. euteiches* culture filtrates subjected to heat treatment (Heated, 100°C, 20 min) or size-based fractionation (100 kDa and 30 kDa molecular weight cut-off membranes). 0% filtrate and 100% unfractionated filtrate serve as negative and positive controls, respectively. Significant differences relative to the 0% filtrate control are indicated (*** p < 0.001; ns: not significant; ANOVA, p = 1.7×10⁻⁶). Points represent individual biological replicates; error bars show standard error of the mean.

To gain insight the biochemical nature of the inducing signal, the *A. euteiches* culture filtrate was subjected to heat treatment and size-based fractionation (Fig. 7E; ANOVA, p = 1.7×10⁻⁶). Heating the filtrate did not significantly reduce its inducing activity compared to the unheated 100% filtrate. Sequential filtration through 100 kDa and 30 kDa molecular weight cut-off membranes revealed that the inducing activity was abolished in the ≤30 kDa fraction, which failed to induce significantly higher DAPG levels than the 0% filtrate control (ns). In contrast, the <100 kDa fraction retained significant inducing activity. Collectively, these results indicate that *A. euteiches* releases heat-stable extracellular compound(s), recovered within the 30–100 kDa fraction, that act as chemical cues to trigger DAPG biosynthesis in antagonistic *Pseudomonas* strains.

## Discussion

The major objective of this study was to investigate the impact of the rhizospheric microbiota on the outcome of the interaction between *M. truncatula* roots and the pathogen *A. euteiches*. To this end, we designed a synthetic community (SynCom) composed of bacteria isolated from the *M. truncatula* rhizosphere, focusing on strains with biological activities related to nutrition, development, or antibiosis. This approach aligns with the growing recognition of SynComs as powerful tools to dissect plant-microbiota interactions, particularly for defining microbial-based solutions to enhance plant resistance to diseases [44].

Infection experiments in gnotobiotic conditions, combined with metabarcoding, metabolomics, and transcriptomics analyses, revealed that infection with *A. euteiches* induced a significant shift in the abundance of specific *Pseudomonas* strains in the rhizosphere. This observation is particularly relevant, as *Pseudomonas* spp. are known to play major roles in plant microbiota, including protection against pathogenic interactions [45]. Metabolomic analyses further demonstrated that, while the SynCom itself did not significantly alter the root metabolite profile, infection with *A. euteiches* led to the detection of new compounds, most notably the *Pseudomonas*-specialized metabolite 2,4-diacetylphloroglucinol (DAPG).

The production of DAPG is highly relevant in the context of legume-*A. euteiches* interactions. Previous studies have shown that the DAPG-producing strain *P. protegens* Pf-5 can efficiently reduce disease symptoms caused by *A. euteiches* in legume plants, and that this protection is dependent on DAPG production [46, 47]. In our study, we confirmed that DAPG-producing strains and purified DAPG inhibit *A. euteiches* growth. Transcriptomic analyses of root tissues further revealed that the plant’s response to the pathogen is strengthened in the presence of DAPG-producing strains, consistent with reports that DAPG can induce defense responses in plants such as *Arabidopsis* [48, 49]. Genome analysis of the two DAPG-producing strains included in the SynCom (A05H and D08H) identified putative biosynthetic gene clusters (BGCs) for additional antimicrobial compounds, including the volatile hydrogen cyanide (HCN) and the lipopeptidic siderophore histicorrugatin [50, 51]. However, neither volatile anti-*Aphanomyces* activity nor iron-dependent inhibition was detected in our assays, strongly suggesting that DAPG is the primary factor mediating the biological activity of these strains against *A. euteiches*.

The enrichment of DAPG-producing *Pseudomonas* during *A. euteiches* infection could be a consequence of DAPG production itself. Recent studies have shown that antimicrobial metabolites such as DAPG, pyoluteorin, and orfamide A can promote the invasion of bacterial communities by *Pseudomonas* strains, due to the susceptibility of other SynCom members to these compounds [52]. Additionally, the production of antimicrobial compounds by roots in response to *A. euteiches* could also inhibit the susceptible bacterial strains. While we did not test the susceptibility of each SynCom member to DAPG or plant metabolites, the significant decrease in bacterial diversity and the increased colonization of root surfaces by these strains observed during *A. euteiches* infection indicates a clear competitive advantage for DAPG-producing strains in this altered rhizospheric environment.

Interestingly, DAPG was not detected in the rhizosphere of healthy plants or in cultures grown in PDB medium, indicating that its production is tightly controlled. At the molecular level, DAPG biosynthesis is regulated by the TetR repressors PhlH and PhlF, as well as post-transcriptionally by the Gac/Rsm pathway [43]. Environmental signals, including carbon sources (e.g., glucose, fructose, mannitol) and root exudate composition, are known to influence DAPG production in various *Pseudomonas* strains [53–56]. Additionally, the plant host genotype can play a role, as demonstrated in wheat, where ancient varieties were more effective at inducing DAPG biosynthetic genes [57]. While we cannot exclude an impact of *M. truncatula* root exudates on DAPG production during infection, our results suggest a major effect of pathogen perception in triggering DAPG biosynthesis. To date, only a few studies have described the inductive or repressive effects of pathogens on DAPG production. For example, the fungal metabolite fusaric acid, produced by *Fusarium oxysporum*, has been shown to inhibit DAPG production and the protective activity of DAPG-producing strains [58, 59]. Although activation of DAPG biosynthetic genes in co-culture plates of a *Pseudomonas* strain with the root pathogen *F. culmorum* has been reported [60], induction of DAPG production upon pathogen perception has not, to our knowledge, been previously described. This novel finding suggests that *A. euteiches* may release specific signals that activate DAPG biosynthesis in rhizospheric *Pseudomonas*. As a first attempt to characterize these signals, we showed that they are heat-resistant and present in a 30 kDa-100 kDa fraction Several studies have been reported on the characterization of mechanisms involved in the recognition of fungi by bacteria and the induction of antifungal mechanisms [61] and it was reported recently that fungal cell wall compounds such as chitin fragments induce antifungal activity in the enteropathogen *Yersinia pseudotuberculosis* [62]. Future work will focus on identifying the specific *A. euteiches*-derived signals that induce DAPG production in *Pseudomonas*.

Our results suggest that induction of anti-oomycete response in rhizospheric bacteria could potentially be a part of a broader mechanism of “extended immunity” which can include the stimulation of microbial defense mechanisms upon the perception of pathogen by rhizobacteria independently of plant-derived signals. The “cry for help” hypothesis posits that plants under biotic stress release compounds that recruit beneficial microbes to suppress pathogens or stimulate defense responses [63]. Our study reveals that perception of the pathogen by rthizospheric bacteria induces a dual defense response including the enrichment of DAPG-producing *Pseudomonas* during *A. euteiches* infection coupled with defense induction leading to the production of antimicrobial metabolites in root exudates (e.g., isoflavonoids [64, 65] and triterpenoids such as betulinic aldehyde [66]). The altered rhizospheric environment during infection may favor the growth of antagonist strains.

From an applied perspective, our findings highlight the potential of DAPG-producing *Pseudomonas* strains as biocontrol agents against *A. euteiches*, a major threat to legume crops. The pathogen-specific induction of DAPG production suggests that these strains could act as “on-demand” protectors, minimizing ecological side effects in the absence of disease pressure and favoring the establishment of suppressive soil condition. Comparative trials with commercial biocontrol strains (e.g., *P. protegens* Pf-5) and field tests under diverse edaphoclimatic conditions will be essential to validate their efficacy. Additionally, the variability in DAPG induction among *M. truncatula* genotypes suggests opportunities for breeding legume varieties that enhance the recruitment of protective microbiota via root exudates, as proposed for wheat and other crops [57]. The design of resilient SynComs— enriched in DAPG-producing strains and resistant to DAPG-mediated collapse—could offer a sustainable alternative to chemical fungicides. However, the ecological trade-offs of DAPG production, such as potential impacts on non-target soil organisms (e.g., mycorrhizal fungi or nitrogen-fixing bacteria), must be carefully evaluated. Furthermore, the stability of DAPG-mediated protection under field conditions, where abiotic factors (e.g., soil pH, moisture) may influence its efficacy, remains to be assessed.

To conclude, this study demonstrates that the rhizosphere microbiota of *M. truncatula* can dynamically respond to *A. euteiches* infection by enriching for DAPG-producing *Pseudomonas* strains, thereby enhancing plant defense. By integrating synthetic community approaches with multi-omics analyses, we provide a framework for dissecting the chemical and genetic underpinnings of microbiome-mediated disease suppression. These insights pave the way for the development of microbiome-based strategies to improve legume resilience, with broader implications for sustainable agriculture.

## Author contributions

MZ, TR and BD conceived the study, GM, AA, MZ, SF, TR, EG, LC and BD analysed the data, AA, MZ, MS, AP, QB performed the research, MZ, TR and BD wrote the paper, TR. and B.D. supervised the study and acquired the funding.

## Acknowledgments

We thank the GeT-Biopuces platform (INSA/TBI/Genotoul Toulouse) member of IBISBA-FR (https://doi.org/10.15454/08BX-VJ91) for performing the libraries and Miseq sequencing.

## Data availability

The datasets generated and analysed during the current study are available in public repositories. Genomics, Transcriptomics, and Metabarcoding: Raw sequencing reads and assembled genomes of the Medicago truncatula root-derived synthetic community (SynCom) under *A. euteiches* infection have been deposited in the National Center for Biotechnology Information (NCBI) BioProject database under accession number PRJNA1422028. Untargeted Metabolomics: The raw metabolomics data have been deposited in the Zenodo repository and are publicly accessible via the Digital Object Identifier (DOI): https://doi.org/10.5281/zenodo.21106495.

## Supporting information

**Supplementary figure 1:**
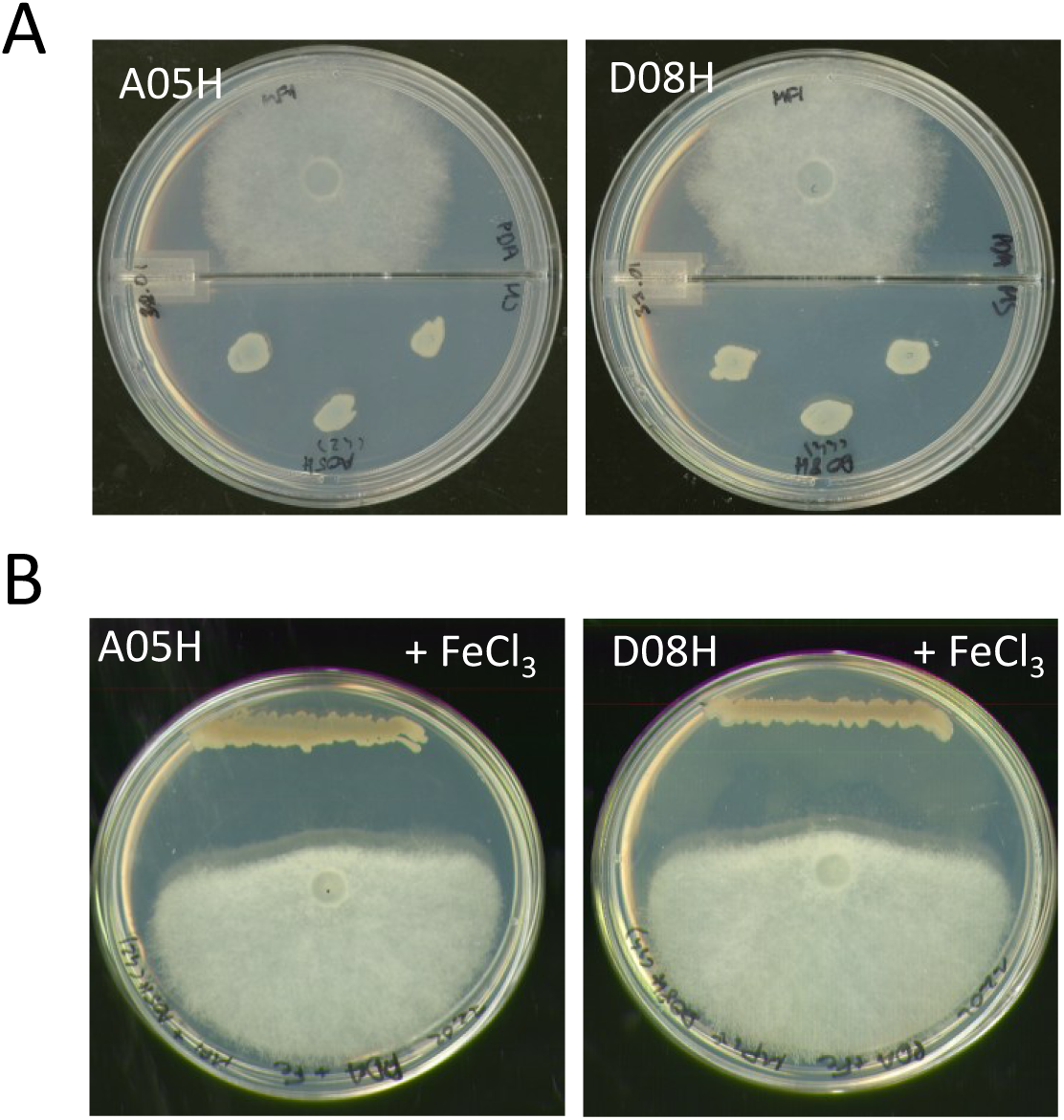
**A.** Co-culture of *A. euteiches* (MF1) with A05H or D08H strains on compartmented Petri dishes, 4 days after inoculation (PDA medium). In the absence of direct contact, Aphanomyces hyphae grow normally, indicating that bacterial volatile compounds are not involved in growth inhibition. **B.** Co-culture of *A. euteiches* with A05H and D08H on iron-complemented PDA, 4 days after inoculation. The growth of *A. euteiches* remains strongly impacted despite the presence of iron in the medium, indicating that siderophore activity is not involved in the antagonistic activity of A05H and D08H.

**Supplementary Table S1**: List of bacterial strains composing the SynCom derived from the

*M. truncatula* rhizosphere, with their functional traits (auxin production, phosphate solubilization, TCP, nitrogen fixation, anti-*Aphanomyces* activity), 16S rRNA sequences, and NCBI best hits.

**Supplementary Table S2**: Relative abundance (%) of bacterial genera detected in the rhizosphere of *M. truncatula* in the SynCom and SynComAphano conditions, and in the T0 inoculum (n = 3 biological replicates per condition).

**Supplementary Table S3**: Untargeted LC-HRMS metabolomic dataset of Medicago truncatula rhizosphere exudates. Each row corresponds to a metabolic feature identified by MS-Dial 5.2 and annotated with MS-Finder. Peak intensities are reported for each biological replicate across the four experimental conditions (Mt, Mt_Ae, Mt_SynC, Mt_Ae_SynC)

**Supplementary Table S4:** Differentially Expressed Genes in roots treated with the SynCom (L2FC_SY), *A. euteiches* (L2FC_AE) and SynCom and *A. euteiches* (L2FC_AESY) relatively to the plant alone. DEGs are genes showing differential expression >1 or <-1 and a FDR <0.001 in at least one condition.

**Supplementary Table S5:** Top induced genes in at least one condition (Log2 FC>4). Only shown are genes with a putative function.

**Supplementary Table S6:** Expression data of gene model predicted to code enzymes involved in flavonoid biosynthesis. Genes were manually annotated upon data and literature mining. The data was used to construct the histograms on the figure 4.

**Supplementary Table S7:** Genomic features and predicted biosynthetic gene clusters (BGCs) of the DAPG-producing strains A05H and D08H. Assembly and annotation statistics were obtained from Flye assemblies annotated with Prokka; BGCs were predicted with antiSMASH (values indicate percentage similarity to the closest reference cluster).

